# Molecular control of dormancy transitions throughout the year in the monoecious cork oak

**DOI:** 10.1101/2024.07.03.601814

**Authors:** Helena Gomes Silva, Rómulo Sobral, Ana Teresa Alhinho, Hugo Ricardo Afonso, Teresa Ribeiro, Patrícia M. A. Silva, Hassan Bousbaa, Leonor Morais-Cecílio, Maria Manuela Ribeiro Costa

## Abstract

Bud dormancy plays a vital role in flowering regulation and fruit production, being highly regulated by endogenous and environmental cues. Deployment of epigenetic modifications and differential gene expression control bud dormancy/break cycles. Information on how these genetic and epigenetic mechanisms are regulated throughout the year is still scarce for temperate trees, such as *Quercus suber*. Here, the expression levels of *QsCENL* and *QsDYL1* during different seasonal cycles of bud development suggest that *QsCENL* may be implicated in the establishment of growth cessation in *Q. suber* and that *QsDYL1* is a good dormancy marker. Moreover, the analysis of the profiles of epigenetic marks and the expression of modifiers, in dormant versus non-dormant bud meristems, indicate that epigenetic regulation is implicated in how bud development progresses in *Q. suber*. The identification of bud specific mechanisms opens new possibilities to understand how trees respond to challenging environmental signals derived from climate change.

## Introduction

In temperate perennial plants, bud dormancy is an adaptive response enabling plants to survive during unbefitting environmental cues such as winter low temperatures (reviewed in Lloret, Badenes & Ríos 2018; Singh *et al*. 2017). Dormancy establishment, maintenance and release are critical moments during the yearly cycle of the plant, influencing the timing of the emergence of reproductive structures (Böhlenius *et al*., 2006; Varkonyi-Gasic *et al*., 2013). The dormant period occurs with a transient interruption of visible growth, which requires mechanisms restricting cell division and expansion. Studies on dormancy suggest that its release is often determined by the exposure to a number of hours of low temperatures (chilling requirement) (Fuchigami and Wisniewski, 1997; Falavigna *et al*., 2019). When dormancy is established, growth is inhibited by intrinsic factors and the bud remains inactive even when subjected to favorable conditions, until the fulfilment of chilling requirements. After that, the growth remains arrested until suitable environmental cues such as of temperature, light and water supply occur, allowing subsequent bud swelling (Anderson *et al*., 2001). The mechanisms by which dormancy is regulated by day length or/and temperature is species dependent (Ruttink *et al*., 2007; Heide and Prestrud, 2005). Phenological evidence in different species has suggested that temperate trees are particularly sensitive to abnormal warm temperatures during autumn, which lead to an out-of-season growth reassumption and blooming. The occurance of flowering in autumn may deplete flower meristems that should otherwise remain dormant inside the buds until spring, with a potential negative impact on the reproductive success of the tree. Therefore, the understanding of the regulatory networks controlling how bud progresses during autumn and winter is of particular importance to provide potential markers of dormancy status in a scenario of future warmer climate.

*Quercus suber* is a temperate Fagaceae tree that inhabits regions with contrasting climatic seasons (mild wet-winters and hot dry-summers) (Pereira, 2007). The environmental parameters required for growth cessation and bud swelling in *Q. suber* remain poorly known (Pinto *et al*., 2011). *Q. suber* presents a complex bud biology, making it difficult to predict the exact timing of bud dormancy release and flower emergence (Sobral *et al*., 2020). In summer, during bud development and before growth cessation, male inflorescence primordia differentiate inside the bud (Sobral *et al*., 2020). Usually during September/October, *Q. suber* bud set and shoot growth cessation occurs, lasting until the end of winter or early spring, when bud swelling occurs simultaneously with the emergence of male flowers. A few weeks later, female inflorescences emerge at the axils of newly formed leaves (Sobral *et al*., 2020). In *Q. suber*, abnormal vegetative growth and male flowering are frequently observed in unusual warm autumns.

There is some information about the molecular mechanisms by which dormancy induction and release occur in buds of temperate perennial species. The better recognized examples of genes controlling bud development are the *DORMANCY ASSOCIATED MADS BOX* (*DAM*) genes and *FLOWERING LOCUS T* (*FT*), with a known involvement during dormancy establishment and release in different species. For instance, in Populus, the orthologs of the Arabidopsis gene *FT*, *PtFT2* controls vegetative growth cessation, bud set and dormancy induction (Hsu *et al*., 2011) while *PtFT1* expression induces early flowering (Böhlenius *et al*., 2006). In the *evergrowing* (*evg*) mutant of peach (*Prunus persica*) the deletion of a locus containing several tandemly duplicated *DAM* genes prevents dormancy induction (Bielenberg *et al*., 2004, 2008; Yamane *et al*., 2008; Li *et al*., 2009; Jiménez *et al*., 2010; Sasaki *et al*., 2011; Wu *et al*., 2012; Saito *et al*., 2013; Hao *et al*., 2015; Howe *et al*., 2015; Li *et al*., 2017; Wu *et al.,* 2017). Moser et al. (2020) showed that silencing *MdDAM1* in apple results in defective autumn growth cessation. *DAM* genes are related to *SHORT VEGETATIVE PHASE* (*SVP*) and *AGAMOUS-LIKE 24* (*AGL24*) of *Arabidopsis thaliana* (Bielenberg *et al*., 2008), where *SVP* responds to cold temperatures to repress (Hartmann *et al*., 2000) and *AGL24* to promote flowering (Michaels *et al*., 2003). *SVP*/*DAM* genes are also involved in the annual dormancy cycle of *Quercus suber* (Sobral *et al*., 2020). The expression of *QsSVP1* and *QsSVP4* gene expression is high during axillary bud set and then steadily decreased until March, coincident with the axillary bud swelling in both adult and juvenile trees (Sobral *et al*., 2020).

The periodic expression of *SVP*-like/*DAM* genes is also thought to be controlled by the increase and/or decrease of epigenetic marks such as trimethylation of histone H3 at lysine 4 (H3K4me3), H3 and H4 acetylation (H3ac, H4ac), trimethylation at lysine 27 of histone H3 (H3K27me3) and DNA methylation in specific genomic regions (Horvath *et al*., 2010; Leida *et al*., 2012; de la Fuente *et al*., 2015; Saito *et al*., 2015; Rothkegel *et al*., 2017). In peach, higher expression of *DAM* genes during the chilling period is correlated with the absence of H3K27me3 marks, while the repression of *DAM* genes during the growing season correlates with an increase in 21-nucleotide small (s)RNAs and noncoding (nc)RNAs, H3K27me3 and CHH methylation (Zhu *et al*., 2020). Recently, the analysis of genome-wide H3K4me3 modifications in dormant pear buds showed an enrichment of the mark around the transcription start site (TSS) of pear *DAM* genes in the beginning of an artificial chilling accumulation treatment period, decreasing until the end of the treatment (Gao *et al*., 2021). A study of the DNA methylome during dormancy in sweet cherry showed that transcriptome reprograming (including a *SVP*-like gene) was preceded by modifications in DNA methylation levels and that these modifications may enhance cold tolerance (Rothkegel *et al*., 2020). Dormancy is also controlled by epigenetic modifications in the sweet chestnut tree (*Castanea sativa*), another perennial Fagaceae species, where the increase and decrease of DNA methylation levels in buds coincide with bud set and bud swelling, respectively (Santamaría *et al*., 2009, 2011).

Other genes may be involved in the establishment and release of dormancy. In poplar, *CENTRORADIALIS*-*LIKE 1* (*CENL1*) delays dormancy release and bud swelling (Ruonala *et al*., 2008; Mohamed *et al*., 2010; Rinne *et al*., 2011; Song and Chen, 2018). On the other hand, *DORMANCY-ASSOCIATED PROTEIN 1* (*DRM1*/*DYL1*) is considered as a dormancy marker whose expression was firstly detected in axillary buds of pea and in Arabidopsis axillary shoots that were forced to enter dormancy (Stafstrom *et al*., 1998; Tatematsu, 2005). *DRM1*/*DYL1* is also upregulated in dormant buds of grapevine (Pacey-Miller *et al*., 2003; Pucker *et al*., 2020; Velappan *et al*., 2022), blueberry (*Vaccinium corymbosum*) (Naik *et al*., 2007), kiwifruit (Wood *et al*., 2013) and sessile oak (*Quercus petraea*) and in dormant seeds of Arabidopsis (Carrera *et al*., 2008).

To better understand the molecular mechanisms that control dormancy processes in *Q. suber*, we characterized the expression of several gene homologs to known regulators of dormancy and epigenetic modulation during bud development in consecutive years. We also investigated whether the deposition of epigenetic marks occurs differently in different types of meristems in dormant and non-dormant buds. This study provides tools that may be used to monitor bud status throughout the seasons, which is of particular importance in the current context of climate change.

## Material and methods

### Plant material

Axillary and apical bud samples were collected monthly from adult and juvenile *Q. suber* trees located at the University of Minho campus (https://www.icampi.uminho.pt/pt/ambiente/green/, *latitude: 41°33 ‘33.8’’ N, longitude: 8°23’54.3’’ W),* throughout four annual cycles (summer 2015 to spring 2019). All samples were taken from the tree crown from 9:00 a.m. to 10:00 a.m. using pruning shears, if not mentioned otherwise. The monthly samples of 2015/2016, 2016/2017, 2017/2018 were constituted by a mixture of approximately twenty randomly collected buds. In 2018/2019, each monthly sampling was made in triplicates, which were used as biological replicates. To analyze circadian gene expression profile, buds were also collected from adult trees at different time points on the course of the day (24 hours, 4 hour interval), in October, January and April.

### RNA extraction and cDNA preparation

Total RNA was obtained using the CTAB/LiCl extraction method (Chang *et al*., 1993) with some modifications (Azevedo *et al*., 2003). Total RNA was treated with DnaseI (Rnase-free) (New England Biolabs) and 0.5-1 ug of RNA was used for cDNA synthesis using the Invitrogen cDNA synthesis kit SuperScript® RT, according to the manufacturer’s instructions.

### Quantitative RT-PCR (qRT-PCR) analysis

Amplification was carried out with SsoFast™ EvaGreen®Supermix (Bio-Rad), 250 nM of each gene-specific primer (Table S1) and 1 μL of cDNA. Quantitative real-time PCR (RT-qPCR) reactions were performed in triplicates on the CFX96 Touch™ Real-Time PCR Detection System (Bio-Rad). After an initial period of 3 min at 95 °C, each of the PCR cycles consisted of a denaturation step of 10 s at 95 °C and an annealing/extension step of 10 s at the temperature optimized for each gene specific primers. With each PCR reaction, a melting curve was obtained to check for amplification specificity and reaction contaminations, by heating the amplification products from 60 °C to 95 °C in 5 s intervals. Primer efficiency was analyzed with CFX Manager™ Software v3.1 (Bio-Rad), using the Livak calculation method for normalized expression (Livak and Schmittgen, 2001). Gene expression analysis was conducted using three technical replicates and normalized with the reference genes *QsPROTEIN PHOSPHATASE 2A SUBUNIT A3* (*QsPP2AA3*) and *ELONGATION FACTOR-1ALPHA* (*QsEF-1α*) (Marum *et al*., 2012).

### Tissue fixation, embedding and sectioning

For morphological and immunolocalization analysis, buds were collected and immediately fixed in 4% (w/v) paraformaldehyde in 1× PBS (phosphate buffered saline: 1x PBS 130 mM NaCl, 7mM Na_2_HPO_4_, 3 mM NaH_2_PO_4_, pH 7.4) under vacuum infiltration followed by overnight incubation in the fixative solution at 4°C. Samples were then dehydrated, cleared and paraffin embedded according to the protocol described by Coen *et al*. (1990) with some modifications: during the clearing procedure, the tissues were incubated 1 hour at room temperature, in each of the following solutions: 75% (v/v) ethanol/25% (v/v) Histo-Clear® (VWR Chemicals), 50% (v/v) ethanol/50% (v/v) Histo-Clear®, 25% (v/v) ethanol/75% (v/v) Histo-Clear®; and incubated three times for 1 hour in 100% (v/v) Histo-Clear®. Tissue sections with 8 µm thickness were obtained using a microtome (SLEE) and mounted in pre-coated poly-L-lysine slides (VWR), deparaffined by heating at 65°C during 15 minutes and washed with Histo-Clear® six times (15 minutes each), and further rehydrated with a grade of ethanol series (100, 80, 70, 50 and 30% (v/v), 5 minutes in each solution) ending with deionized water twice (5 minutes each).

### Immunolocalization of 5-Methylcytosine and Post-translational Histone Modifications

Sections were permeabilized with sodium citrate buffer (10 mM Sodium citrate, 0.05% (v/v) Tween 20, pH 6.0) and by a heat-mediated antigen retrieval treatment, using the microwave at maximum power, for 4 minutes according to Nic-Can *et al*. (2013). After cooling down for 20 minutes, the sections were washed three times in 1x PBST (130 mM NaCl, 7mM Na_2_HPO_4_, 3 mM NaH_2_PO_4_, 0.05% (v/v) Tween). After blocking with 8% (w/v) BSA in 1x PBS for 30 min at 4 °C, the sections were incubated with primary antibodies diluted in 1x PBST: anti-5mC (1:100 dilution, Abcam AB10805), anti-H3K4me3 (1:50 dilution, Abcam AB8580), and anti-H3K18ac (1:100 dilution, Abcam AB1191) at 4 °C during 48 hours in a moist chamber. Negative controls were done by replacing the primary antibody by 1 x PBST. After washing in 1x PBST buffer (3 times, 5 minutes), samples were incubated for three hours at 37 °C, using a solution containing goat polyclonal secondary antibody to mouse or rabbit IgG – H&L conjugated to Alexa Fluor 488 (Abcam AB150113, AB150077), added in a 1:100 dilution. The slides were mounted in EverBrite™ Mounting Medium with 4’, 6-diamidino-2-phenylindole (DAPI, Biotium) and were examined with a Spinning Disk Carl Zeiss Axio Observer Z1 microscope using 10x magnification or with a Plan Apo 63x Ph3-NA1.4 –DIC HCII Oil immersion objective. Images were acquired with an AxioCamMR3 camera controlled by Axiovision 4.8.2.0 software. Zeiss fluorescence filters Set 49 and Set 38 HE were used to excite DAPI, and Alexa Fluor 488, respectively. Z-stacks with 0.2 µm were acquired to allow for the entire nuclei capture. The images of antibody detection and DAPI staining were acquired using the same microscope settings and light exposure times. Evaluation of nuclei fluorescence intensity of z-stack maximum intensity projections was performed using Fiji software (Schindelin *et al*., 2012) after image deconvolution using Huygens Essential (Scientific Volume Imaging B.V., Netherlands). Nuclear staining intensity was evaluated, at least, in 30 nuclei per meristem. Background fluorescence intensity was measured as the average intensity of inter-nuclear areas and divided from the nuclear staining intensity of each nucleus for correction. The intensity measurement of each epigenetic mark was obtained by dividing the labelling intensity of specific signal in each nucleus by the correspondent DAPI intensity, as previously described (Bian *et al*., 2009; Leida *et al*., 2012; Testillano *et al*., 2013; Ingouff *et al*., 2017; Zhai *et al*., 2018). Data shown is representative of at least three independent experiments and the values are represented by arbitrary units.

### Homology-based search and phylogenetic analysis

In order to identify *Q. suber* homologs of known genes involved in dormancy in other species, pea and Arabidopsis DYL (Stafstrom *et al*., 1998; Tatematsu, 2005), poplar CENL1 (Mohamed *et al*., 2010) protein sequences were used as queries for BLASTp searches at NCBI for *Q. suber* (Qs), *V. vinifera* (Vv), *O. sativa* (Os), *Jatropha curcas* (Jc), *M. domestica* (Md), *A. thaliana* (At), *Q. robur* (Qr), *P. sativum* (Ps), *P. persica* (Pp) and *P. trichocharpa* (Pt) homologs. The full-length amino acid sequences of each homolog (Table S2) were aligned with CLUSTALW, using the Gonnet Protein Weight Matrix. The Multiple Alignment Gap Opening penalty to 3 and the Multiple Alignment Gap Extension penalty to 1.8 (Hall, 2013) were used as alignment parameters. Phylogenetic analyses were performed with MEGA7.0 software program (Kumar *et al*., 2016) using the Jones-Taylor-Thornton (JTT) correction model, inferred by the Maximum-likelihood algorithm. The resulting trees were constructed based upon 1000 bootstrap replicates.

### Statistical analysis

The raw data were imported into GraphPad Prism V6.0 software to calculate descriptive statistic parameters such as means and standard deviations.

Differences in the labelling fluorescent intensity/DAPI intensity between tissues and/or developmental phases were tested through two-way ANOVA followed by Tukey’s multiple comparison test, at a 5% significance level.

### Meteorological support data

Air temperature and accumulated cold hours (<7.2 °C) were determined based on the monthly weather report bulletin (Instituto Português do Mar e da Atmosfera-IPMA) with the data monitored at the nearest meteorological station (Braga - Merelim, 41.57586944,-8.45110833, 65) from University of Minho campus (41.56236104, −8.39401066).

## Results

### The complex nature of active growth and dormancy periods in *Quercus suber*

*Quercus suber* trees have meristematic tissues and organ primordia enclosed in terminal and axillary buds that cease activity and enter dormancy during autumn (Fig S1 A), possibly in response to shorter days and/or lower temperatures, similarly to what has been described for other Quercus species (Derory *et al*., 2006). The buds contain vegetative (Fig 1A, red arrows) or both vegetative and reproductive meristems (Fig 1B, highlighted in red boxes) and stay dormant throughout winter, probably due to the persistency of low temperatures. Usually, in late February/March, the increase in day length and/or rising temperatures signals the end of the dormancy period and induces bud swelling (Fig 1C). Male flowers with well-defined anthers (Fig 1D) emerge in the axils of the swollen axillary buds (Fig S1 B). In April/May, male flowers complete their development and new shoots emerge containing young leaves and female flowers (Fig S1 C). A new axillary bud emerges in the axils of the leaves that are not flanked by a female flower spike (Fig 1E and FIG S1 D). In the buds collected in June/July (Fig S1), the differentiation of male catkin primordia (Fig 1F and 1G, red boxes) adjacent to the apical meristem (Fig 1F, red arrow) was observed.

**Figure 1.**
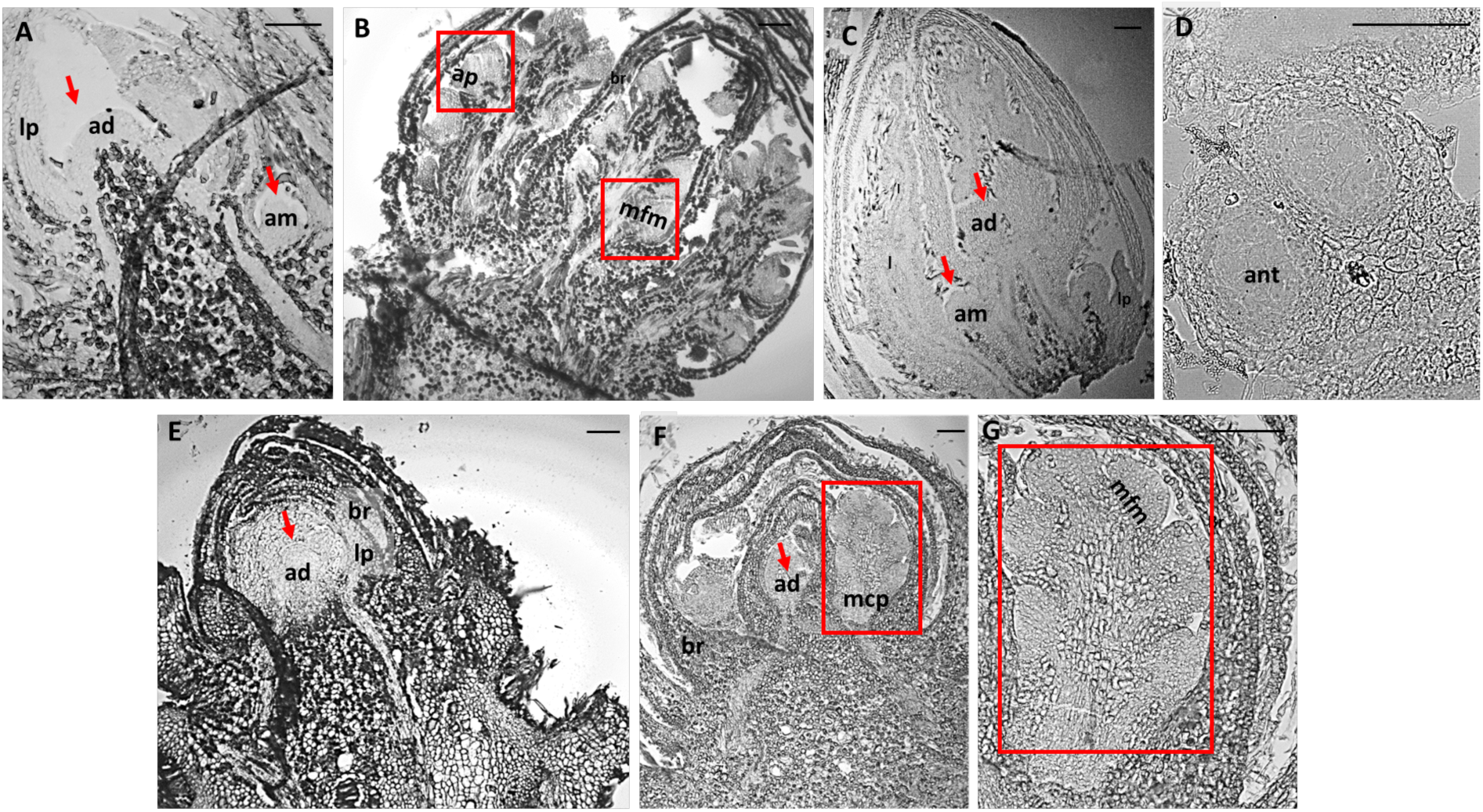
Histological sections of vegetative and reproductive stages of *Quercus suber* buds. **A)** Longitudinal section of a vegetative bud during bud set in September containing a shoot apical meristem and axillary meristem protected by scales. **B)** Longitudinal section of a reproductive bud in September containing male catkin primordia with male flower meristem and anther primordia. **C)** Longitudinal section of an April swollen bud containing axillary meristems in the axils of leaf primordia. **D)** Transversal section of a male flower, that emerged in the axil of an April swollen bud, showing developed anthers. **E)** Axillary bud that developed in spring in the axils of new leaves containing a shoot apical meristem. **F)** Actively growing buds in June containing an apical meristem and inflorescence primordia (square) with **G)** male flower meristems. ad: apical meristem, am: axillary meristem, ap: anther primordia, br: bracts, ip: inflorescence primordia, lp: leaf primordia, mcp: male catkin primordia and mfm: male flower meristem. Scale: 100 µM.

The vegetative and reproductive development of *Q. suber* shows other particularities that are frequently observed in the field. In some years with unusual warm temperatures in autumn or early winter, premature bud swelling occurs with the emergence of male flowers (but rarely female flowers). In spring, many buds fail to swell, and the timing of bud swelling fluctuates, between and within trees.

### *QsCENL* and *QsDYL1* might be involved in dormancy

It has been previously suggested that *CENL* and *DYL1* may suppress bud growth in other species (Soeda, 2004; Liu *et al*., 2007; Ruonala *et al*., 2008; Mohamed *et al*., 2010; Rinne *et al*., 2011; Lee *et al*., 2013; Footitt *et al*., 2017) but it is not known if the function is conserved in Fagaceae. Homologous sequences of CENL and DYL1 family members were retrieved from the *Q. suber* genome by homology-based search. CENL belongs to the FT family, together with FT TERMINAL FLOWER 1 (TFL1), TWIN SISTER OF FT (TSF) and MOTHER OF FT (MFT) (Wickland and Hanzawa, 2015).

The phylogenetic tree was divided into distinct clades, each one associated to a different sub-family (Fig 2A). Two homologs belong to the TFL1 clade (QsTFL1 and QsCENL), another to the FT clade (QsFT) and the fourth to the MFT clade (QsMFT) (Fig 2A).

**Figure 2.**
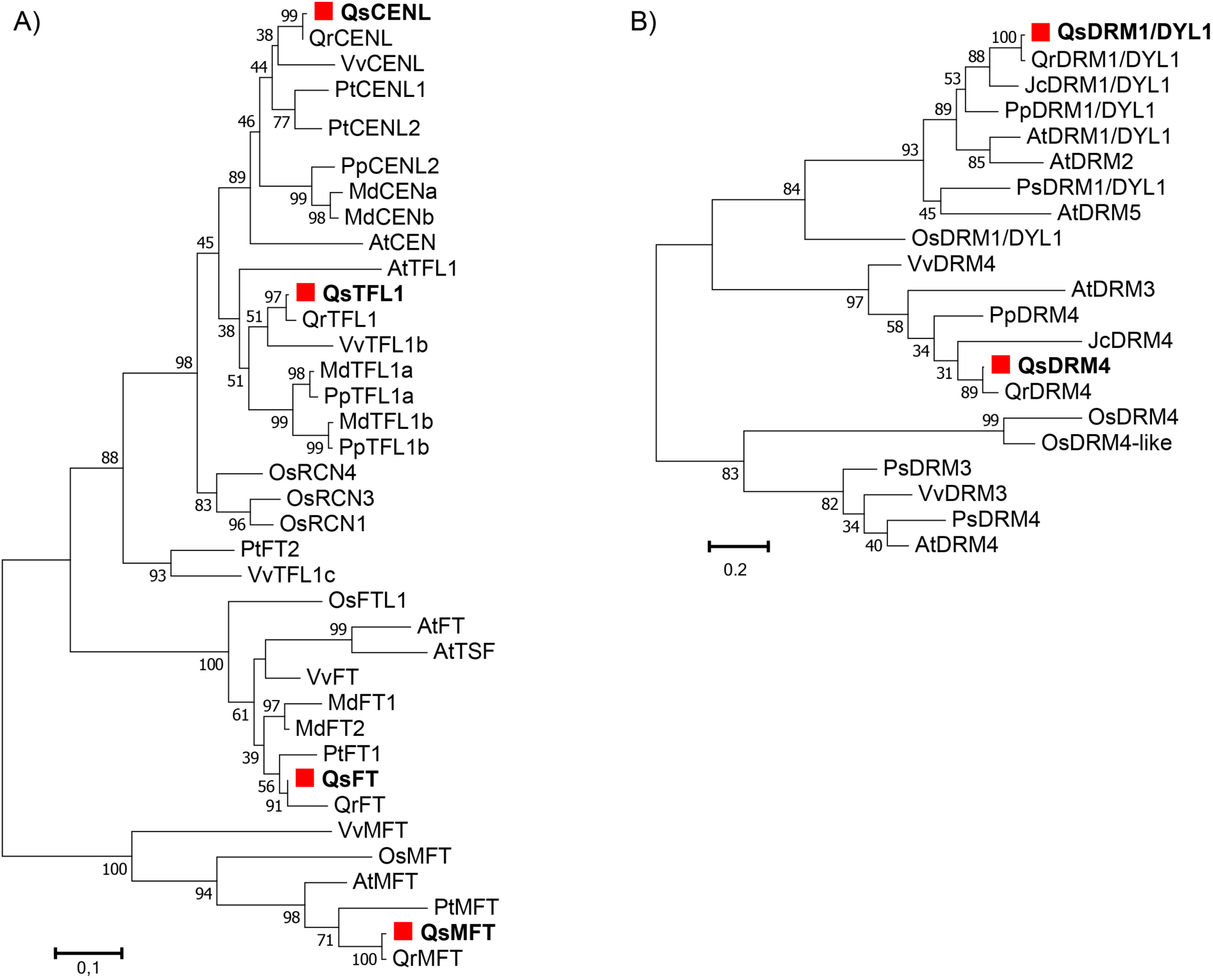
Identification of A) QsCENL and B) QsDYL1 by phylogenetic analysis. The *Q. suber* proteins are shown in bold and highlighted with a red square. The phylogenetic relationship of each family proteins was inferred using the Maximum-likelihood method. The percentage of replicate trees in which the associated taxa clustered together in the bootstrap test (1000 replicates) is shown next to the branches. The evolutionary distances were computed using the JTT matrix-based method and are in the units of the number of amino acid substitutions per site. Protein accession numbers used in the phylogenetic analysis are presented in Table S2.

Two homologous proteins of DYL1 were found in *Q. suber* but only one belongs to the DRM1/DYL1 clade (Fig 2B). The phylogenetic tree suggested that QsDYL1 is the closest homolog to PsDYL1/DRM1 and AtDRM1, considered as good dormancy markers in axillary buds of pea and in Arabidopsis seeds, respectively (Stafstrom *et al*., 1998; Tatematsu, 2005).

The transcription levels of *CENL* and *DYL1* were evaluated during the bud growth-dormancy cycle in four consecutive years. *QsCENL* expression shows a conserved profile in all the years under study: it is higher during bud set (September/October), then decreases and remains at low levels until March (Fig 3). However, *QsCENL* was upregulated in the swollen buds (in April) in two of the periods (2015/2016; 2018/2019) (Fig 3). To distinguish the involvement of *QsCENL* in dormancy vs flowering development, its expression was also analyzed in buds collected from *Q. suber* juvenile trees (not competent to flower) in October, January and March. The expression pattern was the same as the one observed in adult trees, higher expression in October and low expression throughout the following months (Fig S2). To analyze if the *QsCENL* expression profile is affected by the circadian rhythm, buds were also collected from adult trees at different time points on the course of the day (24 hours, 4 hour interval), in October, January and April. The expression of *QsCENL* was higher in October bud set regardless of the time of day or night (Fig S3).

**Figure 3.**
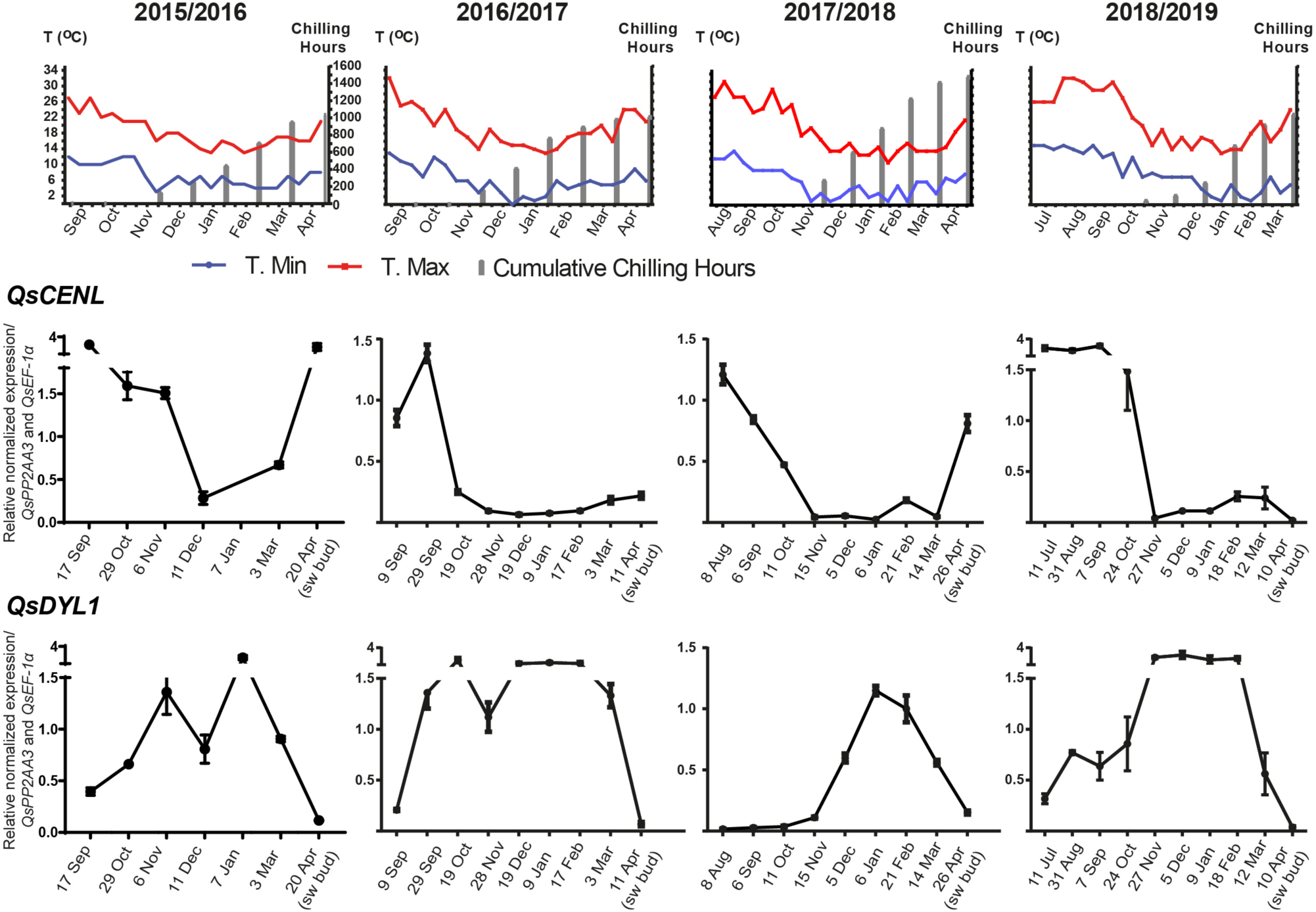
Expression of *QsCENL* and *QsDYL1* during bud growth-dormancy transitions. The transcript accumulation in the buds were analyzed by RT-qPCR and the normalized values are represented in each graph. Corresponding annual growth cycle are shown in the upper part of each graph, along with graph showing the average of minimum and maximum weekly temperature for each month and the accumulation of chilling hours (<7.2 °C). The name of the gene is shown in the upper left side of the four equivalent graphs. Expression levels are relative to *QsPP2AA3* and *QsEF-1α*. Sampling dates are shown in x-axis.

In adult trees, *QsDYL1* expression increases after bud set, usually in September/October (Fig 3). Then, the expression decreases until the emergence of swollen buds in April (Fig 3). In juvenile trees, high levels of *QsDYL* expression were found in October, then, the expression decreases until the end of dormancy time (March), similarly to what was observed in adult trees (Fig S2). However, levels of *QsDYL* expression had significant variation between the years of study. Similarly, bud samples collected at different time points during the day in October and January had significant variation regarding *QsDYL* transcript levels (Fig S3). Nevertheless, the same trend was observed: higher expression during bud dormancy (January) (Fig S3 and Fig 3).

### The expression of DNA methyltransferases and histone acetyltransferases is switched on and off throughout bud development

A likely epigenetic control of bud dormancy in Fagaceae was first reported in *C. sativa* with emphasis in DNA methylation and histone acetylation (Santamaría *et al*., 2009; 2011). To assess the role of DNA methylation and histone acetylation during bud development, the expression of genes coding for enzymes that catalyze the deposition of these epigenetic marks was analyzed. The *Q. suber* epigenetic modifiers were phylogenetically identified in a previous study (Silva *et al*., 2020). Here, we studied gene expression of five DNA methyltransferases (*QsMET1*, *QsDRM2*, *QsDNMT2*, *QsCMT3* and *QsCMT1*) and three histone acetyltransferases (*QsHAF1*, *QsHAM1* and *QsHAC1*) during bud development (Fig 4 and S3). *QsCMT1* transcript accumulation in the buds was very low (Fig S4). An increase in *QsMET1* expression was observed prior to April bud swelling (Fig 4), while *QsDNMT2* expression generally increased after October and decreased in April (Fig 4). *QsCMT3* was highly expressed during late stages of bud formation, when the buds were still growing (July to August/September), but the expression decreased gradually from bud set (September/October) to bud dormancy (Fig 4). The expression increased again from February to April, being highly expressed in swollen buds (April) (Fig 4). The expression of *QsHAC1* was higher during growth cessation and dormancy when compared to bud swelling (Fig 4). *QsHAF1* and *QsHAM1* expression profiles were not conserved in the different years (Fig S4).

**Figure 4.**
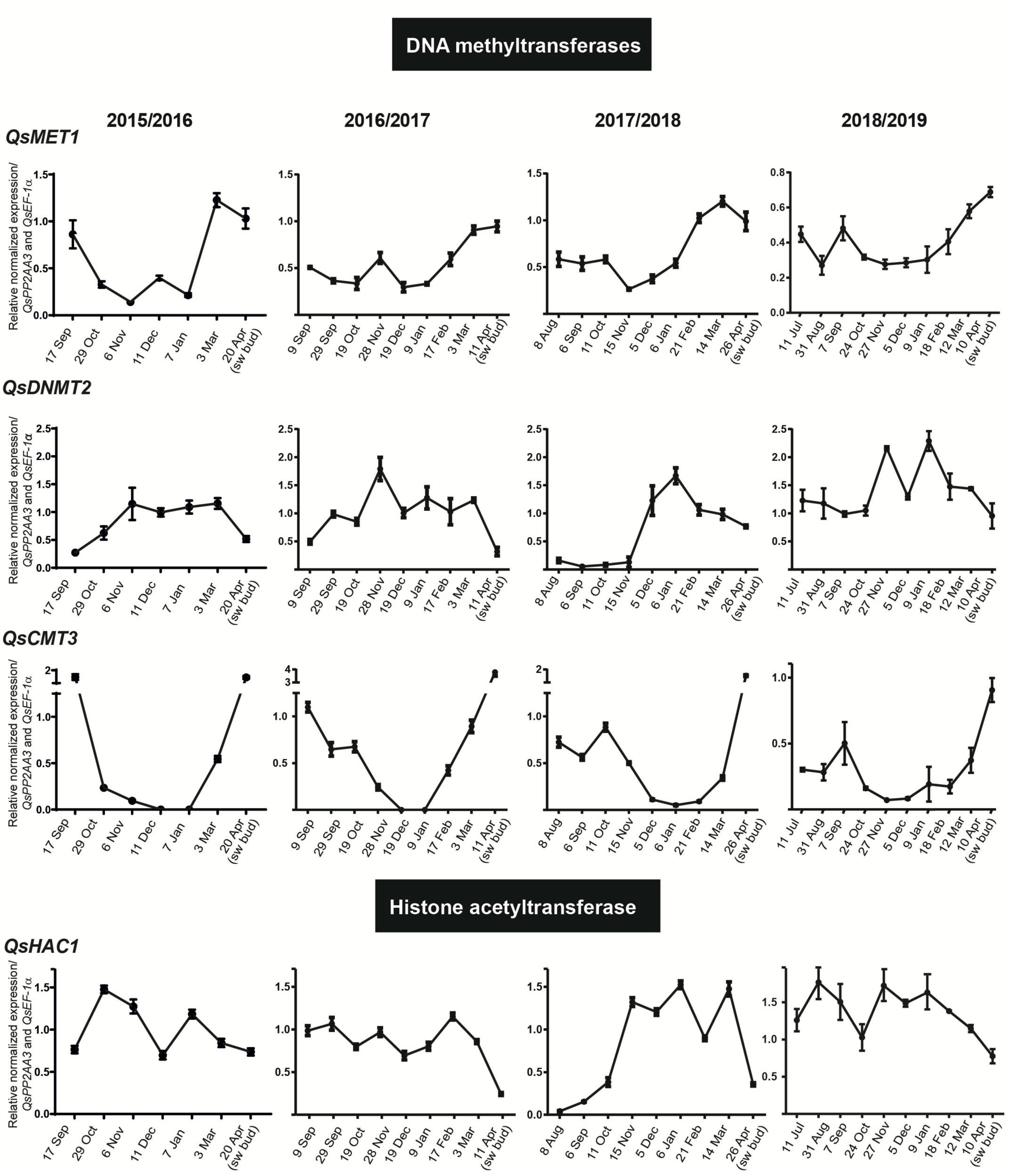
Expression of *Quercus suber* epigenetic regulators during bud growth-dormancy transitions. DNA methyltransferases *QsMET1*, *QsDNMT2*, and *QsCMT3* and Histone acetyltransferases *QsHAC1* expression patterns were studied by RT-qPCR in buds and the normalized values are represented in each graph. The corresponding annual growth cycle are shown in the upper part of each graph. The name of the gene is shown in the upper left side of the four equivalent graphs. Expression levels are relative to *QsPP2AA3* and *QsEF-1α*. Sampling dates are shown in x-axis.

To infer if *QsCMT3* up- or down-regulation is associated exclusively with bud dormancy or bud growth instead of reproductive development, its expression was measured in vegetative buds of juvenile trees during two different years. *QsCMT3* was downregulated in both years during dormancy establishment (from October to December) and upregulated during dormancy release (March) (Fig S2) in a similar manner to adult trees. The buds collected from adult trees at different time points of the day were also used to analyze the circadian regulation of *QsCMT3*. The results showed that the gene is always highly expressed in the swollen opened buds (April) and less expressed in the months where bud set is occurring or dormancy is already established (October and January), regardless of the time of the day at which the samples were collected (Fig S3).

### Differential distribution of epigenetic marks in dormant and non-dormant meristems

To get insights into how chromatin modifications may affect dormancy, the distribution of epigenetic marks in buds at different developmental stages was analysed. Immunolocalization of the repressive mark 5-methylcytosine (5mC) and active marks such as acetylation of histone H3 at lysine 18 (H3K18ac) and trimethylation of histone H3 at lysine 4 (H3K4me3) (Fig 5) were performed in buds containing both reproductive and/or vegetative meristems or tissues before growth cessation in summer, in dormant winter buds, and during bud swelling in the spring.

**Figure 5.**
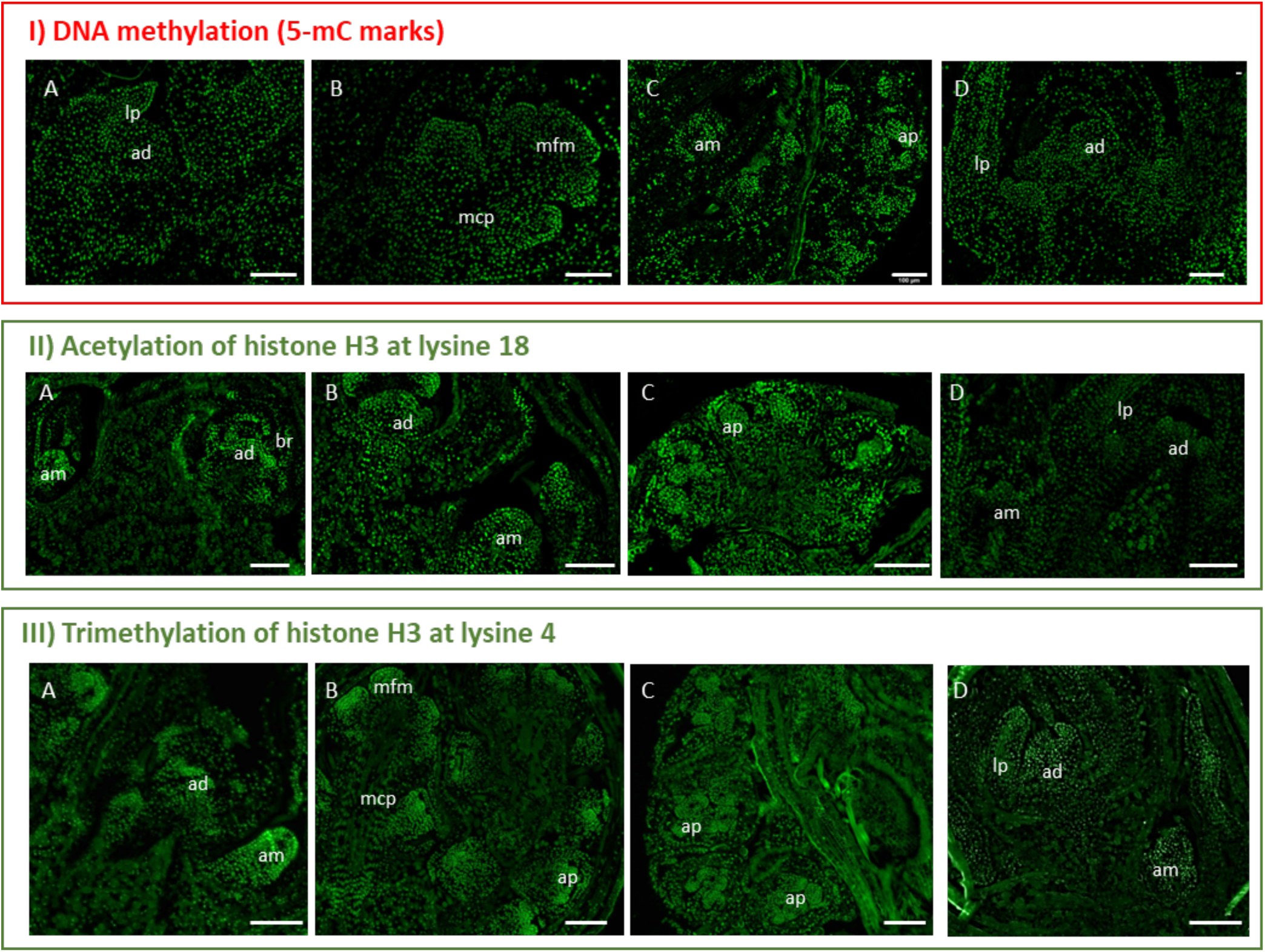
I) DNA methylation (5-mC marks), II) Acetylation of histone H3 at lysine 18 and Trimethylation of histone H3 at lysine 4 in axillary buds of *Quercus suber*. Immunolocalization of epigenetic marks (green signal) in longitudinal sections of **(A-B)** late stages of bud formation, **(C)** bud dormancy and **(D)** bud swelling. Scale bars: 100 μm. ad: apical meristem, am: axillary meristem, ap: anther primordia, br: bracts, lp: leaf primordia, mcp: male catkin primordia and mfm: male flower meristem.

During late stages of bud formation in the summer, the three epigenetic marks were detected in both apical and axillary meristems (Fig 5A - I, II, III). The marks were also observed in early stages of male catkin primordia and in the flower meristems found in the already differentiated male catkins (Fig 5B – I, III). Throughout dormancy, the marks were likewise observed in both vegetative and reproductive primordia (Fig 5C - I, II, III), in male flowers and also in the apical and axillary meristems within the swollen bud (Fig 5D - I, II, III).

The intensity level of each epigenetic mark was evaluated in apical and axillary meristems, as well as in male flower meristems, during late stages of bud formation (summer), bud dormancy (winter) and anters during bud swelling period (spring) (Fig 6, Fig 7 and Fig 8). A higher intensity of 5-mC signal was detected in male flower meristems and in the apical meristems of dormant buds, when compared to the corresponding structures of summer and spring (Fig 6). H3K18 acetylation and DNA methylation in the apical meristem showed an opposite pattern regarding the transition from late stages of bud formation to bud dormancy (Fig 6 and Fig 7). In both apical and axillary meristems, H3K18ac signal intensity is higher during late stages of bud formation (Fig 7D and Fig 7F). On other hand, male flower meristems (Fig 7B) had lower levels of H3K18ac in comparison to the terminal (Fig 7D) and axillary meristems (Fig 7F). H3K18ac appeared to be similarly represented in male flower meristems throughout the dormancy-growth cycle (Fig 7B, Fig 7H, Fig 7N). Tri-methylation of H3K4 is weaker in male flower meristems of dormant buds (Fig 8H) when compared with the ones found during late stages of bud formation (Fig 8B) and during male flower emergence (Fig 8B). During bud swelling period, H3K4me3 is significantly higher in male flower meristems (Fig 8N) than in axillary meristems (Fig 8R).

**Figure 6.**
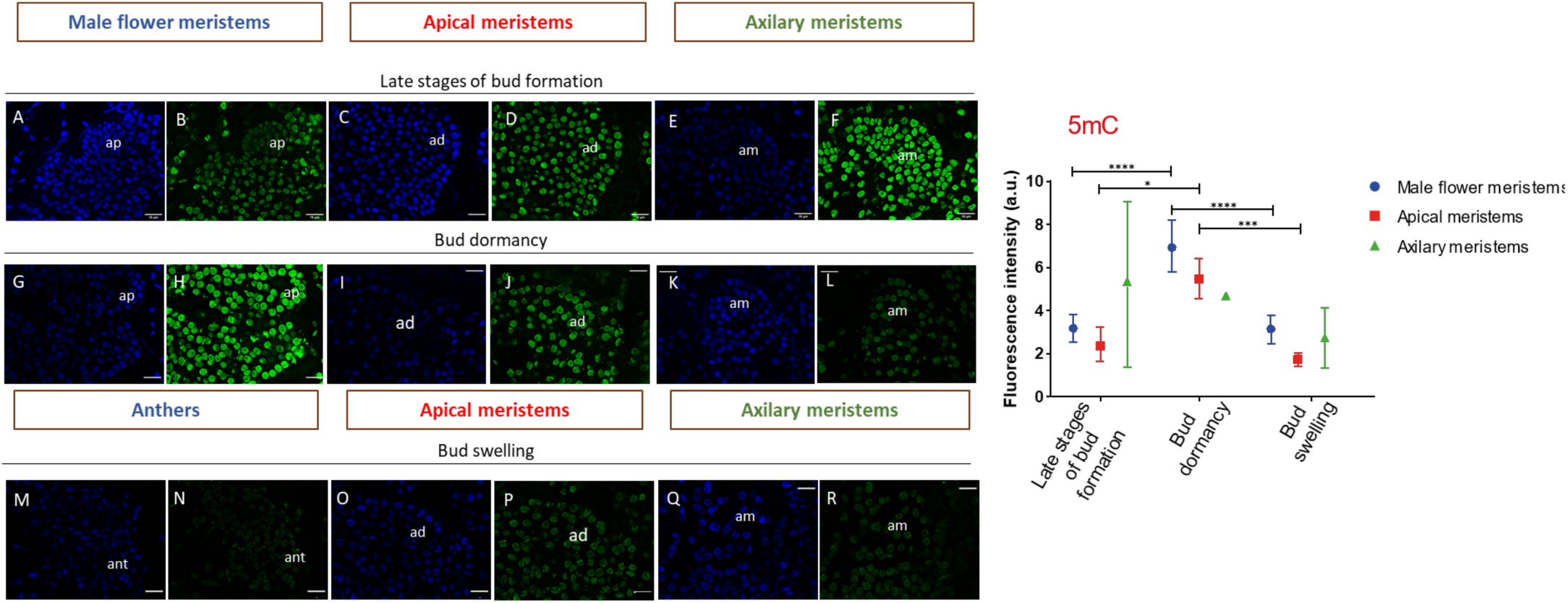
Quantification of DNA methylation in the different type of meristems inside *Quercus suber* buds, during late stages of bud formation in summer, in winter dormant buds and in the spring swollen buds. **Right panel:** The represented values are the biological replicate mean of the calculated average labelling intensity of each nuclei relative to the mean of DAPI labelling in each type of tissue, as measured in the deconvoluted maximum projection images. Sampling stages are shown in x-axis. Error bars indicate standard deviation (SD) of at least three biological replicates. a.u. arbitrary units. Male flower meristems are represented with blue circles, apical meristems with red squares and axillary meristems with green triangle. Statistically significant pairwise differences are indicated with asterisks (**** P value < 0.0001), according to two-way ANOVA. **Left panel** - Microscopy images used to fluorescence intensity quantification**: (A)** Late stages of bud formation (summer) with male flower meristems showing DNA (blue) and **(B)** H3K18ac signal (green). **(C)** Summer buds with apical meristems showing DNA (blue) and **(D)** H3K18ac signal (green). **(E)** Summer buds with axillary meristems showing DNA (blue) and **(F)** H3K18ac signal (green). **(G)** Winter dormant buds with male flower meristems showing DNA (blue) and **(H)** H3K18ac signal (green). **(I)** Winter dormant buds with apical meristems showing DNA (blue) and **(J)** H3K18ac signal (green). **(K)** Winter dormant buds with axillary meristems showing DNA (blue) and **(L)** H3K18ac signal (green). **(M)** Emerged male anthers of spring showing DNA (blue) and **(N)** H3K18ac signal (green). **(O)** Swollen buds with apical meristems showing DNA (blue) and (**P)** H3K18ac signal (green). **(Q)** Swollen buds with axilary meristems showing DNA (blue) and **(R)** H3K18ac signal (green). ad: apical meristem, am: axillary meristem, ant: anthers, ap: anther primordia and mfm: male flower meristem. Scale bars: 15 μm.

**Figure 7.**
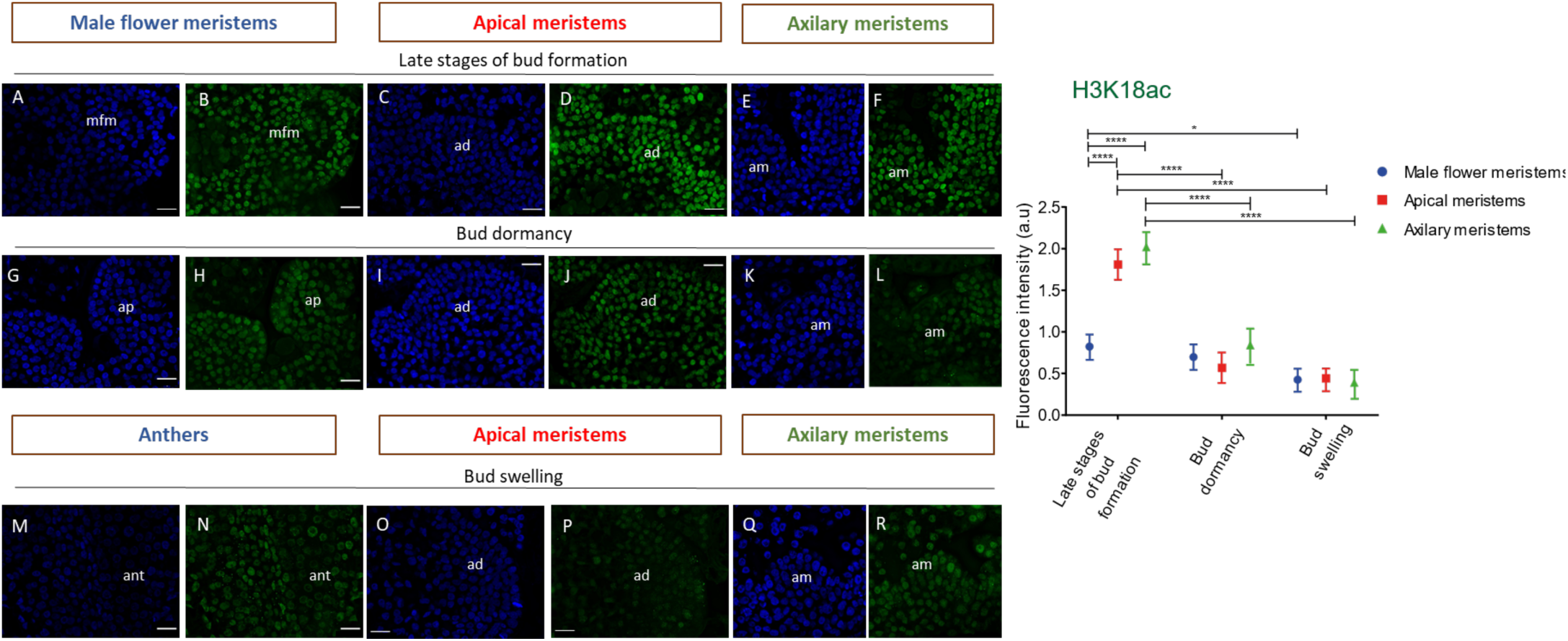
Quantification of histone H3 acetylation at lysine 18 in the different type of meristems inside *Quercus suber* buds, during late stages of bud formation in summer, in winter dormant buds and in the spring swollen buds. **Right panel:** The represented values are the biological replicate mean of the calculated average labelling intensity of each nuclei relative to the mean of DAPI labelling in each type of tissue, as measured in the deconvoluted maximum projection images. Sampling stages are shown in x-axis. Error bars indicate standard deviation (SD) of at least three biological replicates. a.u. arbitrary units. Male flower meristems are represented with blue circles, apical meristems with red squares and axillary meristems with green triangle. Statistically significant pairwise differences are indicated with asterisks (**** P value < 0.0001), according to two-way ANOVA. **Left panel** - Microscopy images used to fluorescence intensity quantification**: (A)** Late stages of bud formation (summer) with male flower meristems showing DNA (blue) and **(B)** H3K18ac signal (green). **(C)** Summer buds with apical meristems showing DNA (blue) and **(D)** H3K18ac signal (green). **(E)** Summer buds with axillary meristems showing DNA (blue) and **(F)** H3K18ac signal (green). **(G)** Winter dormant buds with male flower meristems showing DNA (blue) and **(H)** H3K18ac signal (green). **(I)** Winter dormant buds with apical meristems showing DNA (blue) and **(J)** H3K18ac signal (green). **(K)** Winter dormant buds with axillary meristems showing DNA (blue) and **(L)** H3K18ac signal (green). **(M)** Emerged male anthers of spring showing DNA (blue) and **(N)** H3K18ac signal (green). **(O)** Swollen buds with apical meristems showing DNA (blue) and (**P)** H3K18ac signal (green). **(Q)** Swollen buds with axilary meristems showing DNA (blue) and **(R)** H3K18ac signal (green). ad: apical meristem, am: axillary meristem, ant: anthers, ap: anther primordia and mfm: male flower meristem. Scale bars: 15 μm.

**Figure 8.**
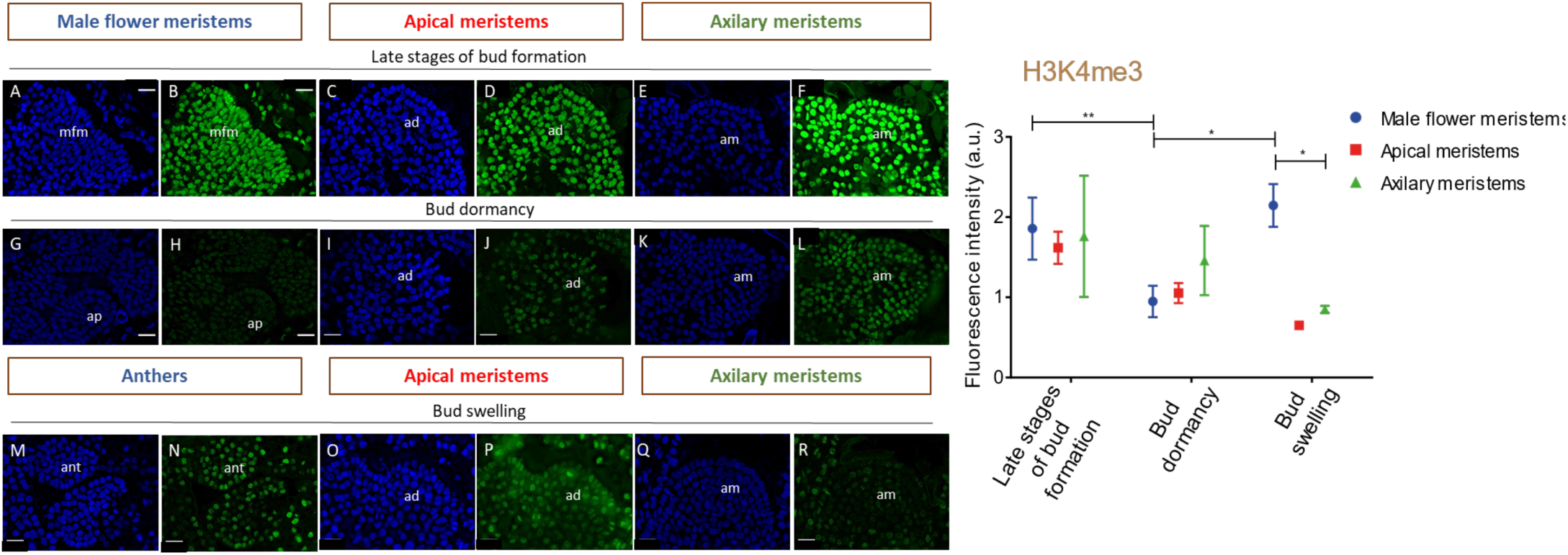
Quantification of the histone H3 trimethylation at lysine 4 in the different type of meristems inside *Quercus suber* buds, during late stages of bud formation in summer, in winter dormant buds and in the spring swollen buds. **Right panel:** The represented values are the biological replicate mean of the calculated average labelling intensity of each nuclei relative to the mean of DAPI labelling in each type of tissue, as measured in the deconvoluted maximum projection images. Sampling stages are shown in x-axis. Error bars indicate standard deviation (SD) of at least three biological replicates. a.u. arbitrary units. Male flower meristems are represented with blue circles, apical meristems with red squares and axillary meristems with green triangle. Statistically significant pairwise differences are indicated with asterisks (**** P value < 0.0001), according to two-way ANOVA. **Left panel** - Microscopy images used to fluorescence intensity quantification: **(A)** Late stages of bud formation (summer) with male flower meristems showing DNA (blue) and **(B)** H3K4me3 signal (green). **(C)** Summer buds with apical meristems showing DNA (blue) and **(D)** H3K4me3 signal (green). **(E)** Summer buds with axillary meristems showing DNA (blue) and **(F)** H3K4me3 signal (green). **(G)** Winter dormant buds with male flower meristems showing DNA (blue) and **(H)** H3K4me3 signal (green). **(I)** Winter dormant buds with apical meristems showing DNA (blue) and (**J)** H3K4me3 signal (green). **(K)** Winter dormant buds with axillary meristems showing DNA (blue) and (**L)** H3K4me3 signal (green). **(M)** Emerged male anthers of spring showing DNA (blue) and **(N)** H3K4me3 signal (green). **(O)** Swollen buds with apical meristems showing DNA (blue) and **(P)** H3K4me3 signa l(green). **(Q)** Swollen buds with axilary meristems showing DNA (blue) and **(R)** H3K4me3 signal (green). ad: apical meristem, am: axillary meristem, ant: anthers, ap: anther primordia and mfm: male flower meristem. Scale bars: 15 μm.

## Discussion

Dormancy transitions are crucial environmental dependent phenophases in perennial plants, since they affect the appropriate time of flower development and consequently fruit production. The identification of the *DAM*/*SVP* genes as promoters of dormancy in peach was one of the major findings regarding molecular regulation of dormancy transitions and, since then, their involvement in dormancy has been further studied in several other perennial plants such as leafy spurge (Horvath *et al*., 2008), apple (Mimida *et al*., 2015), Japanese pear (Ubi *et al*., 2010; Saito *et al*., 2013), tea plant (Hao *et al*., 2017), kiwifruit (Wu *et al*., 2012), Japanese apricot (Yamane *et al*., 2008; Sasaki *et al*., 2011) and in poplar (Singh *et al*., 2018, 2019). The molecular mechanisms that regulate the induction, maintenance and release of bud dormancy remains poorly understood in *Q. suber*. In this species, *QsSVP1* and *QsSVP4* might be involved in the establishment of bud dormancy due to their transcript accumulation (higher during bud set and lower during dormancy and release) and to the ability to delay the flowering time in transgenic Arabidopsis plants (Sobral *et al*., 2020). Other *Q. suber* genes possibly involved in dormancy regulation were studied in this work.

In poplar, *PtCENL1* down-regulation during growth cessation (Ruonala *et al*., 2008), is essential for bud release (Mohamed *et al*., 2010). *PtCENL1* is also upregulated in spring when axillary meristem identity is established (Mohamed *et al*., 2010). In poplar, *PtCENL1* and *PtCENL2* regulate flowering but have an additional role in maintaining inflorescence identity (Mohamed *et al*., 2010). *QsCENL* expression is also higher before and during growth cessation (September/October), decreasing during dormancy (Fig 3), both in adults and juvenile trees, suggesting a role in growth cessation. However, in *Q. suber*, in two of the four studied periods, *QsCENL* was also upregulated during bud swelling (Fig 3), which may be indicative of the presence of new inflorescences meristems inside the buds (unknown at the time of bud collection).

*QsDYL1* expression appears to increase after growth cessation in September/October, it is maintained during during the dormancy period (usually between September/October and February), and it decreases around bud release (Fig 3). Interestingly, upregulation of *QsDYL1* is partially coincident with the downregulation of *QsCENL* (Fig 3). *DYL1* expression was considered a good marker for bud and seed dormancy in other species (Pacey-Miller *et al*., 2003; Tatematsu, 2005; Naik *et al*., 2007; Carrera *et al*., 2008; Ueno *et al*., 2013; Wood *et al*., 2013). Pucker et al. (2020) proposed that *V. vinifera* July buds are already on their way to endodormancy, based on the increasing expression of the *DYL1* homologue (*VviDRM1*). In *Q. suber*, *QsDYL1* may be also a good marker for bud dormancy.

Epigenetic control seems to play an essential role in bud developmental transitions. In this work, we evaluated the expression of several genes coding for enzymes that deposit epigenetic marks. The expression of *QsHAC1* is higher during growth cessation and dormancy when compared to bud swelling (Fig 4). In Arabidopsis, *HAC1* is involved in flowering time regulation by activating the upstream pathways involved in the down-regulation of the main repressor of flowering gene *FLOWERING LOCUS C* (*FLC*) (Deng *et al*., 2007) and is also more expressed during seed dormancy (Footitt *et al*., 2015). If the role of *QsHAC1* in the acetylation of histones is indeed conserved, and consequently associated with active chromatin, perhaps QsHAC1 may target genes that are required to be active in stages in which growth is arrested. *QsMET1* expression is always higher in March, prior to bud swelling (Fig 4), even when *QsDYL* had a higher expression (Fig 4). The upregulation of *QsMET1* seems to precede the swollen bud stage, suggesting that this gene may be involved in sensing bud release signals. In fact, in the vegetative SAM of peach trees, the accumulation of *PpMET1* transcripts was associated with the resumption of growth and with the beginning of floral transition (Giannino *et al*., 2003). *QsCMT3* expression decreases from late stages of bud formation until bud dormancy and then increased again during bud release (Fig 4). This pattern of expression between dormant and non-dormant tissues is completely opposite of *QsDYL1* (Fig 3 and Fig 4). Here we suggest that the expression levels of *QsCMT3* together with *QsDYL1* may help to predict the relative rate of dormancy/growth between samples. CMT3 is essential to methylate cytosines in daughter strands during DNA replication (Bartee *et al*., 2001; Du *et al*., 2012). Due to its crucial action during cell division, higher *QsCMT3* expression is suggestive of higher growth rate, while higher *QsDYL* may be indicative of higher dormancy state. In summary, bud samples with higher *QsCMT3* expression and lower *QsDYL1* expression might contain more actively growing tissues than samples with higher *QsDYL1* expression and lower *QsCMT3* expression. However, it remains unknown if the variation of *QsCMT3* expression observed during the year is specific to bud growth-dormancy transitions or if it is likewise differentially regulated in other tissues in which the temporary growth cessation occurs.

Epigenetic regulation controls key developmental genes during the dormancy-growth cycle. Repressive (DNA methylation and H3K27me3) and active (H3K4me3 and H3ac) epigenetic marks have been shown to be present in the promoter regions of transcription factors such as *SVP-* and *FT*-like genes (kumar *et al*., 2009; Horvath *et al*., 2010; Leida *et al*., 2012; de la Fuente *et al*., 2015; Saito *et al*., 2015; Rothkegel *et al*., 2017; Zhu *et al*., 2020; Gao *et al*., 2021). The immunolocalization of epigenetic marks allowed us to examine changes in 5-methylcytosine DNA and histone modifications at the cellular level in different tissues inside the buds. Different intensity levels of DNA methylation were observed in dormant buds versus actively growing buds, in both apical and male flower meristem nuclei. Fluorescence signal is higher for both meristems in dormant buds (Fig 6). The higher DNA methylation levels in the nuclei of apical meristem in dormant buds is in agreement with what was previously reported in poplar meristems (Conde *et al*., 2013; Le Gac *et al*., 2018) and in chestnut (Santamaría *et al*., 2009). The decrease in DNA methylation during dormancy release was also found in other tissues such as potato tubers and pepper seeds (Law and Suttle, 2002; Portis *et al*., 2004). Our results suggest that the cessation or activation of the apical meristems and male flower primordia growth might be related with some *de novo* methylation or demethylation events. The increase in DNA methylation in dormant buds could be triggered by the *de novo* DNA methyltransferase *QsDRM2* (Fig 6 and Fig S4), repressing large chromatin regions. In contrast, chromatin regions might be more accessible to transcription during bud growth resumption to induce cell division and meristematic activity. So, a higher level of gene activation marks such as H3K4me3 and H3K18ac in male flower primordia, during summer and spring was expected. In fact, in male flower primordia, the levels of tri-methylation of H3K4 had a completely opposite pattern to the 5mC labelling, whereas the acetylation of H3K18 remained constant throughout the year (Fig 7 and Fig 8). The expression of histone acetyltransferases like *QsHAC1, QsHAM1* and *QsHAF1* may play a role in the maintenance of euchromatin in some *loci* in male flower meristems during winter. During later stages of bud formation, H3K18ac accumulation in apical and axillary buds is higher (Fig 7). The reduction in H3K18ac during dormancy might be related with the repression of growth genes, enabling the dormant apical meristem to remain unresponsive in case of exposure to growth-promoting conditions during winter. This was not observed in male flowers (Fig 7). Perhaps, genes involved in male flower development may not be properly inactivated during the dormant period since, in some years, unusual hot temperatures trigger male flowers emergence in autumn. The biological replicates of axillary meristems analysed showed quite different values of 5mC and H3K4me3 fluorescence levels (Fig 6 and Fig 8), which reflect the heterogeinity of these meristems that can originate either a vegetative or reproductive bud

The global intensity of specific histone marks and DNA methylation during the dormancy cycle in *Q. suber* differs between each type of meristems and/or each dormancy stage. Likewise, the expression levels of the enzymes that deposit these marks varies according to the dormancy state, such as *QsCMT3*. Many of the genes discussed in this work appear to have an association with specific dormancy stages and to be part of some molecular regulatory mechanisms that may cooperate to establish, maintain, and release bud dormancy in *Q. suber*. This work provides new insights into the molecular mechanisms that may regulate dormancy transitions in this species and may help to monitor bud status throughout the seasons with particular importance in the context of the current climate change.

