## Supplementary data for "Molecular control of dormancy transitions throughout the year in the monoecious cork oak"

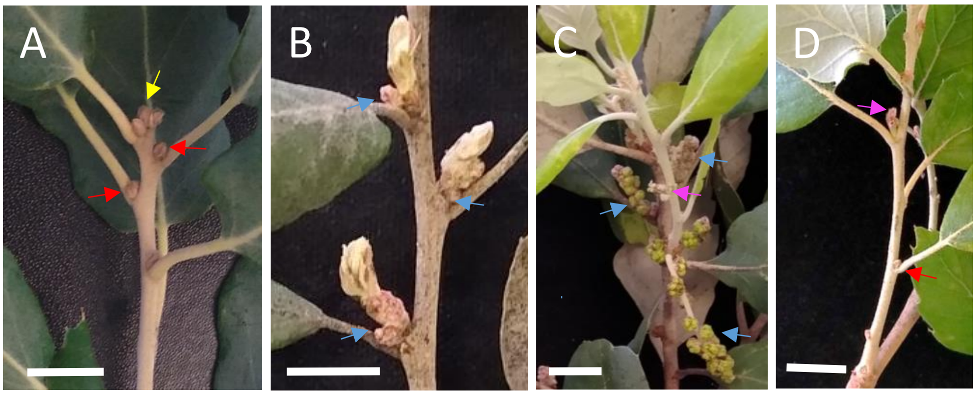

**Supplementary Figure 1. *Quercus suber* buds throughout the year. A)** *Q. suber* branch displaying apical (yellow arrow) and axillary (red arrows) buds during autumn September/October; **B)** After bud swelling in February/March, male inflorescences (blue arrows) emerge in the axils of the axillary buds; **C)** The buds burst and give rise to new branches with female flowers (pink arrow) in the axils in the new leaves (leaf was removed to allow the visualization of the female flower); D) In May, new axillary buds (red arrow) emerge in the axils of the new leaves that are not flanked by a female flower spike (pink arrow). Scale: 1 cm.

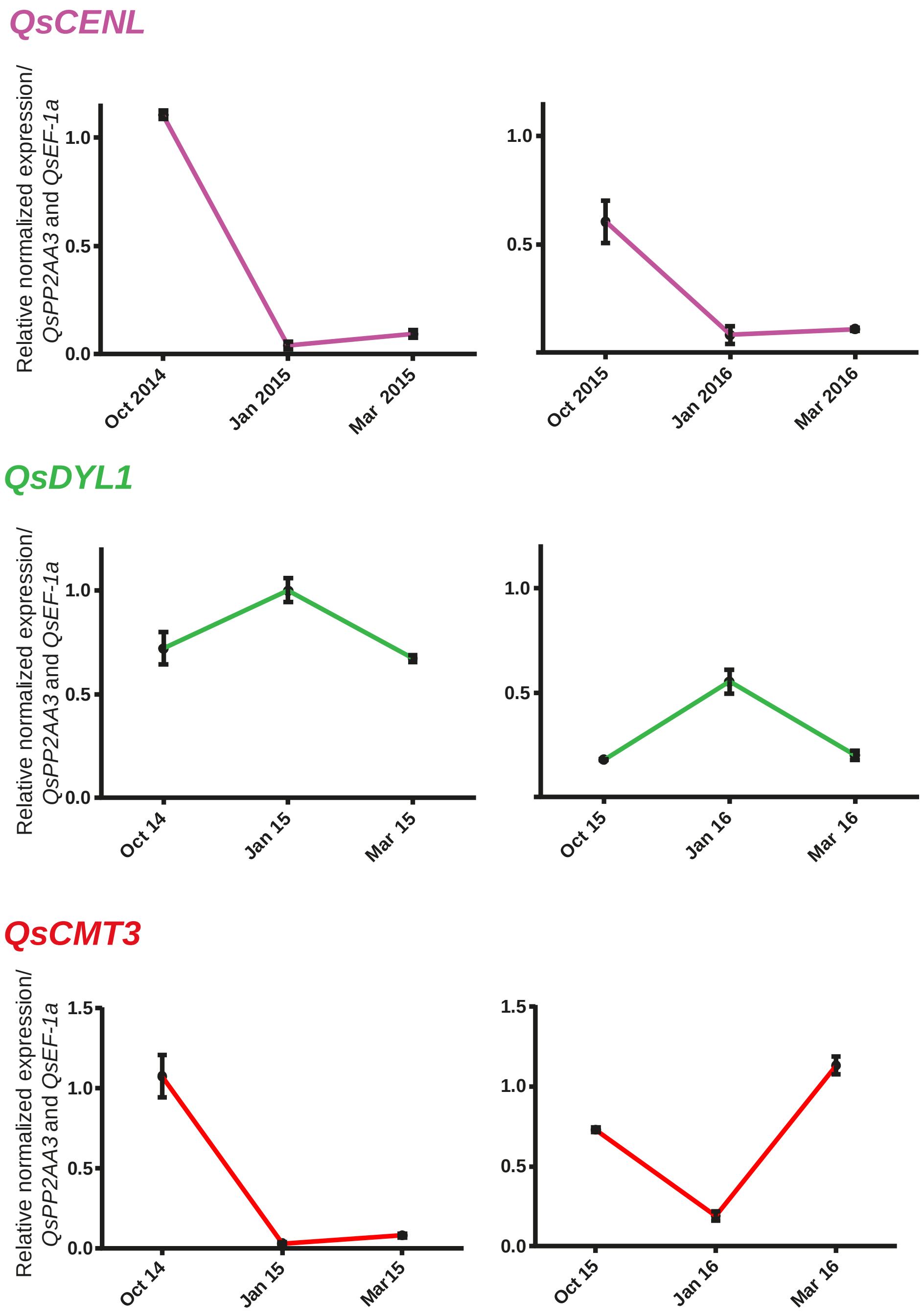

**Supplementary Figure 2. Expression of *QsCENL*, *QsDYL1* and *QsCMT3* in the axillary buds of juvenile *Quercus suber* trees**. October, January and March samples were chosen to represent each stage of bud development (bud set, bud dormancy and bud release, respectively). Two annual growth cycles were studied. The name of the gene is shown in the upper left side of the equivalent graphs. Normalized expression levels are relative to *QsPP2AA3* and *QsEF-1α*. Each value represents expression mean of mixture of buds ± SD of three technical replicates. Sampling dates are shown in x-axis.

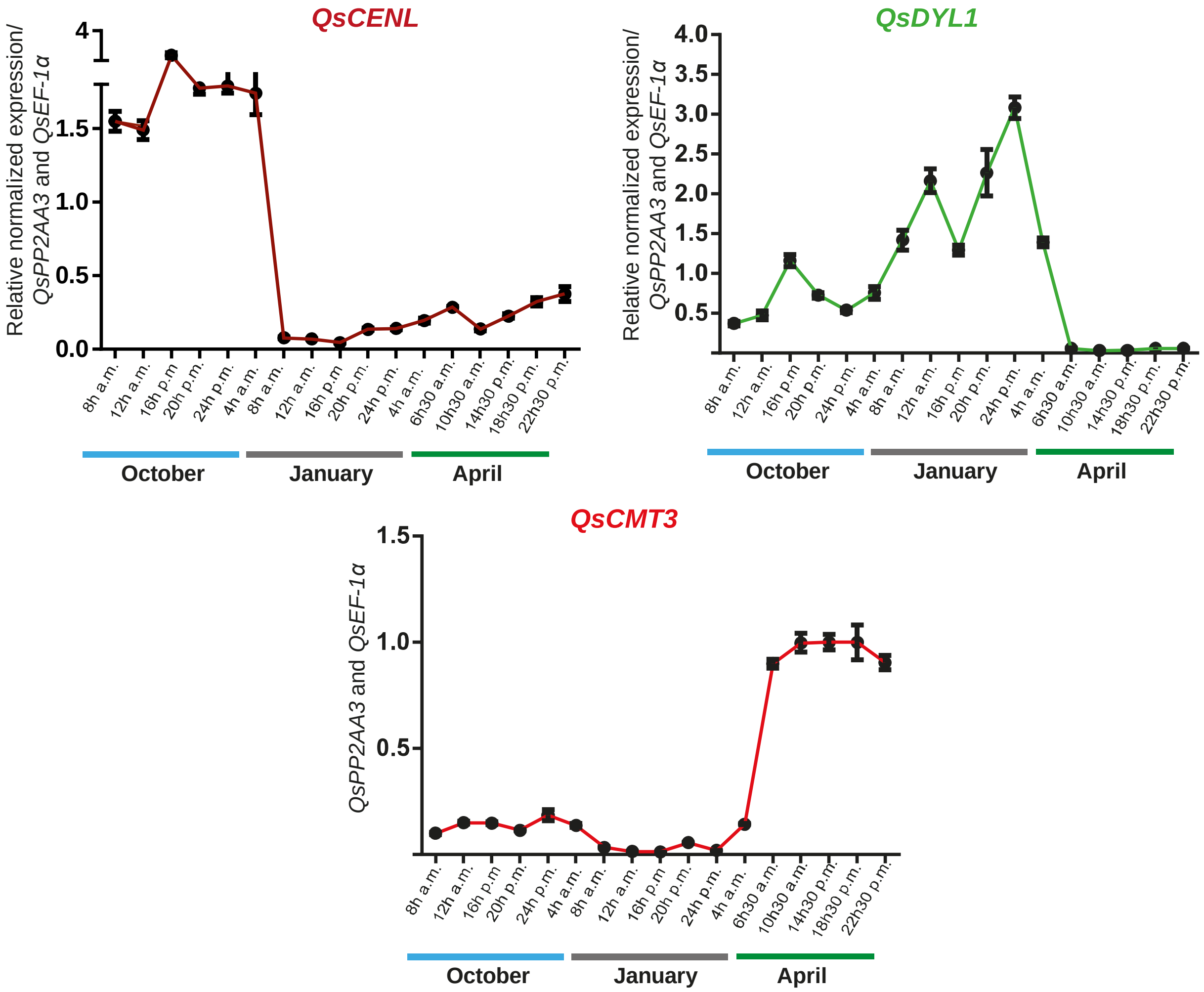

**Supplementary Figure 3. Expression of *QsCENL*, *QsDYL* and *QsCMT3* at different times of the day, under different seasonal conditions**. Buds collected in the same day of October, January and April were collected with four hours intervals since sun rise. RT-qPCR analysis shown in the graphs show the normalized expression relative to *QsPP2AA3* and *QsEF-1α*. The name of the gene is shown in the upper part of each graph. Each value represents expression mean of mixture of buds ± SD of three technical replicates for the first two reproductive seasons. Sampling hours are shown in x-axis.

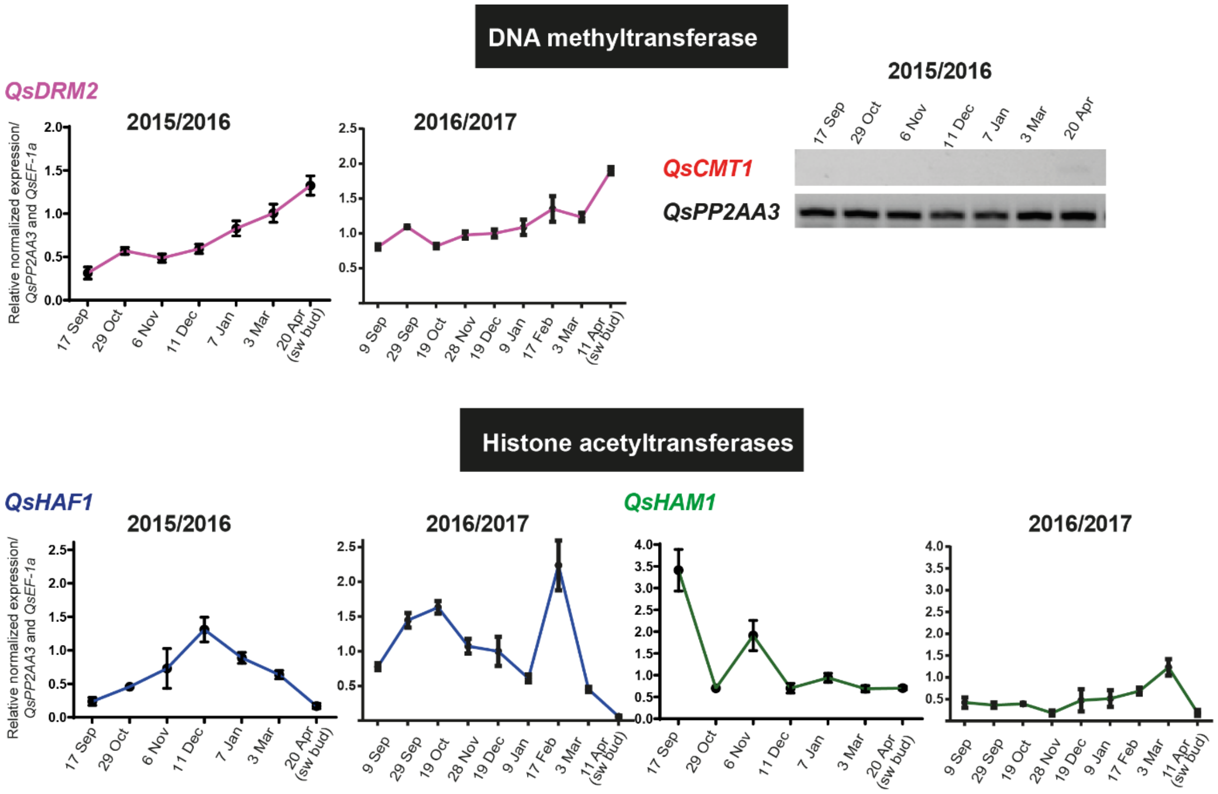

**Supplementary Figure 4. Expression of other *Quercus suber* epigenetic regulators during growth-dormancy transitions.** DNA methyltransferases *QsDRM2* and Histone acetyltransferases *QsHAF1* and *QsHAM1* expression were analysed by RT-qPCR and the normalized values are represented in each graph. The corresponding annual growth cycles are shown in the upper part of the graphs. The name of the gene is shown in the upper left side of each graphs. Normalized expression levels are relative to *QsPP2AA3* and *QsEF-1α*. The DNA methyltransferase *QsCMT1* expression was measured by semi-quantitative PCR, as shown by gel electrophoresis image.

**Supplementary Table S1 – List of Primers**

| Amplicon | Direction | sequence 5'-3' |
|---|---|---|
| *QsCMT3* | Forward | ATGCTGCTGAAGAACACACTCC |
| | Reverse | GCTCGAGCCTTGGGCTATG |
| *QsCMT1* | Forward | CAGCAACTGACGGTTCTTCAC |
| | Reverse | ATTGCCACTTCCTGGTGATGTT |
| *QsMET1* | Forward | TCAACCGCTTCCTTGAGTTT |
| | Reverse | GCTTTCACCCTGAACAGGAC |
| *QsDNMT2* | Forward | CCCCATTGAAAGAGCAGTGT |
| | Reverse | TATGCGTCATTGGATGGAAA |
| *QsDRM2* | Forward | CAAAAGGTGATGCCTGGTCT |
| | Reverse | TAACGATGGGCCAACTTCTC |
| *QsDYL1* | Forward | GATGCTCGCTCTTGGTCTG |
| | Reverse | TATTCCGGTGACACCAACAA |
| *QsHAC1* | Forward | TTCTTGCGACATGCTAATCG |
| | Reverse | AGGAACACAAACAGGGCAAC |
| *QsHAM1* | Forward | ATCCAAAGGTCTTGGATCGTC |
| | Reverse | CCAGGAAGAATCCCCAGTTT |
| *QsHAF1* | Forward | CAAAAGGGTGCCAGAGAGAG |
| | Reverse | CGGTCTTTAAGCGTCTCCAC |
| *QsCMT2.1* | Forward | TTTGCATCTGATTTCGCTTG |
| | Reverse | CCTTCTGAAACCCTGAGACG |
| *QsCMT2.2* | Forward | ATGGATTTGACAGAGCAAAAAAC |
| | Reverse | CATCCTCAGACAAAGATGTTGC |
| *QsDRM3* | Forward | TGGATTCCTCGAATTCAAATG |
| | Reverse | GAGGGTAAGAATCCCATTCCA |
| *QsGCN5* | Forward | GAACCAGTTGATGCACGAGA |
| | Reverse | GCATTGGCAAACATCCTTTT |
| *QsPP2AA3* | Forward | GGGTTCCCAACATCAAGTTC |
| | Reverse | TGACCTGATCACTTGACTGC |
| *QsEF-1α* | Forward | CATCATGAACCACCCCGGTCA |
| | Reverse | CCCAGCATCTCCGTTCTTCA |

**Supplementary Table S2 – List of Protein accessions**

| DYL | | | |
|---|---|---|---|
| **Protein ID** | **Protein name** | **Protein ID** | **Protein name** |
| XP 023929655.1 | QsDRM1/DYL1 | XP 020416550.1 | PpDRM4 |
| OCV3 prime rep c51526 | QrDRM1/DYL1 | XP 012068831.1 | JcDRM4 |
| NP 001295697.1 | JcDRM1/DYL1 | XP 023905393.1 | QsDRM4 |
| XP 007212248.1 | PpDRM1/DYL1 | OCV4 rep c10592 | QrDRM4 |
| At1g28330 | AtDRM1/DYL1 | LOC Os08g35190.3 | OsDRM4 |
| At2g33830 | AtDRM2 | LOC Os09g26620.4 | OsDRM4-like |
| AAB84193.1 | PsDRM1/DYL1 | AAM62421.1 | PsDRM3 |
| At5g44300 | AtDRM5 | XP 002283180.1 | VvDRM3 |
| LOC Os11g44810.3 | OsDRM1/DYL1 | AAM62422.1 | PsDRM4 |
| RVW52172.1 | VvDRM4 | At1g56220 | AtDRM4 |
| At1g54070 | AtDRM3 | | |

| CENL | | | |
|---|---|---|---|
| **Protein ID** | **Protein name** | **Protein ID** | **Protein name** |
| XP 023880417.1 | QsCENL | LOC Os11g05470.1 | OsRCN1 |
| OCV4 c28298 | QrCENL | XP_002321903.2 | PtFT2 |
| NP_001267929.1 | VvCENL | NP_001267933.1 | VvTFL1c |
| Q6TXM3 | PtCENL1 | LOC Os01g11940.1 | OsFTL1 |
| XP 002312811.1 | PtCENL2 | At1g65480 | AtFT |
| XP 007206006.1 | PpCENL2 | AT4G20370.1 | AtTSF |
| NP_001280813.1 | MdCENa | ABF56526.1 | VvFT |
| NP_001280940.1 | MdCENb | ABF84010.1 | MdFT1 |
| AT2G27550.1 | AtCEN | NP_001280810.1 | MdFT2 |
| At5g03840 | AtTFL1 | XP_002316173.1 | PtFT1 |
| XP 023905546.1 | QsTFL1 | XP 023899320.1 | QsFT |
| Loc 111210 Tr 1 2 | QrTFL1 | OCV4 c10192 | QrFT |
| NP_001267931.1 | VvTFL1b | NP_001267935.1 | VvMFT |
| NP_001280887.1 | MdTFL1a | LOC Os06g30370.1 | OsMFT |
| BAD10962.1 | PpTFL1a | At1g18100 | AtMFT |
| NP_001280794.1 | MdTFL1b | XP_002321507.1 | PtMFT |
| NP_001289244.1 | PpTFL1b | XP 023927178.1 | QsMFT |
| LOC Os04g33570.1 | OsRCN4 | Loc 125140 Tr 2 2 | QrMFT |
| LOC Os12g05590.1 | OsRCN3 | | |
